# A stress sensor IRE1α is required for bacterial exotoxin-induced inflammasome activation in tissue-resident macrophages

**DOI:** 10.1101/2023.07.06.547940

**Authors:** Izumi Sasaki, Yuri Fukuda-Ohta, Chihiro Nakai, Naoko Wakaki-Nishiyama, Chizuyo Okamoto, Takashi Orimo, Daisuke Okuzaki, Shuhei Morita, Shiori Kaji, Yuki Furuta, Hiroaki Hemmi, Takashi Kato, Asumi Yamamoto, Takashi Tanaka, Katsuaki Hoshino, Shinji Fukuda, Kensuke Miyake, Etsushi Kuroda, Ken J. Ishii, Takao Iwawaki, Koichi Furukawa, Tsuneyasu Kaisho

**Affiliations:** Department of Immunology, Institute of Advanced Medicine, Wakayama Medical University, Wakayama 641-8509, Japan; Laboratory for Protein Conformation Diseases, RIKEN Center for Brain Science, Wako, Saitama 351-0198, Japan; Genome Information Research Center, Research Institute for Microbial Diseases, Osaka University, Suita, Osaka 565-0871, Japan; First Department of Medicine, Wakayama Medical University, Wakayama 641-8509, Japan; Second Department of Internal Medicine, Wakayama Medical University, Wakayama 641-8509, Japan; Department of Thoracic and Cardiovascular Surgery, Wakayama Medical University, Wakayama 641-8509, Japan; Faculty of Veterinary Medicine, Okayama University of Science, Imabari, Ehime 794-8555, Japan; Laboratory for Developmental Genetics, RIKEN Center for Integrative Medical Science, Yokohama, Kanagawa 230-0045, Japan; Department of Immunology, Faculty of Medicine, Kagawa University, Miki, Kagawa 761-0793, Japan; Institute for Advanced Biosciences, Keio University, Tsuruoka, Yamagata 997-0052, Japan; Gut Environmental Design Group, Kanagawa Institute of Industrial Science and Technology, Kawasaki, Kanagawa 210-0821, Japan; Transborder Medical Research Center, University of Tsukuba, Tsukuba, Ibaraki 305-8575, Japan; Laboratory for Regenerative Microbiology, Juntendo University Graduate School of Medicine, Tokyo 113-8421, Japan; Division of Innate Immunity, Department of Microbiology and Immunology, Institute of Medical Science, University of Tokyo, Tokyo 108-8639, Japan; Department of Immunology, School of Medicine, Hyogo Medical University, Nishinomiya, Hyogo 663-8501, Japan; Division of Vaccine Science, Department of Microbiology and Immunology, The Institute of Medical Science, University of Tokyo, Tokyo 108-8639, Japan; Division of Cell Medicine, Department of Life Science, Medical Research Institute, Kanazawa Medical University, Uchinada, Ishikawa 920-0293, Japan; Department of Biomedical Sciences, Chubu University College of Life and Health Sciences, Kasugai, Aichi 487-8501, Japan

## Abstract

Cholera toxin (CT), a bacterial exotoxin composed of one A subunit (CTA) and five B subunits (CTB), functions as an immune adjuvant. CTB can induce production of interleukin-1β (IL-1β), a proinflammatory cytokine, in synergy with a lipopolysaccharide (LPS), from resident peritoneal macrophages (RPMs) through the pyrin and NLRP3 inflammasomes. However, how CTB or CT activates these inflammasomes in the macrophages has been unclear. Here, we clarified the roles of IRE1α, an endoplasmic reticulum (ER) stress sensor, in CT-induced IL-1β production from RPMs. In RPMs, CTB is incorporated into ER and induced ER stress responses, depending on GM1, a cell membrane ganglioside. IRE1α-deficient RPMs showed a significant impairment of CT- or CTB-induced IL-1β production, indicating that IRE1α was required for CT- or CTB-induced IL-1β production from RPMs. This study first demonstrates the critical roles of IRE1α in activation of both NLRP3 and pyrin inflammasomes in tissue-resident macrophages.

**One sentence summary:** IRE1α is required for NLRP3 and pyrin-mediated IL-1β production

## INTRODUCTION

Certain types of bacteria produce exotoxins, which not only act on intestinal epithelial cells to induce digestive symptoms such as diarrhea but also activate immune cells to function as an immune adjuvant. *Vibrio cholerae*, a gram-negative bacteria, produce cholera toxin (CT). CT is comprised of one A subunit (CTA) and five B subunits (CTB). It also functions as a strong immune adjuvant that can induce Th17 responses, induction of antibody-producing cells, and production of proinflammatory cytokines (*1–6*). CTB can induce production of interleukin-1β (IL-1β), a proinflammatory cytokine, from *in vitro* cultured murine bone marrow derived macrophages (BMMs) in synergy with O111:B4-derived lipopolysaccharides (LPS O111:B4) that can bind to CTB (*7*). In this induction, CTB guides entry of LPS into the cell and internalized LPS activates non-canonical caspases to produce IL-1β (*7*). In murine resident peritoneal macrophages (RPMs), however, CTB, incorporated with ganglioside GM1, can induce IL-1β production in synergy with O55:B5-derived LPS (LPS O55:B5) that fails to bind to CTB. This indicates that CTB does not function as a mere chaperone for LPS, but is directly involved in IL-1β production (*8, 9*). This IL-1β production in RPMs involves activation of NLRP3 and pyrin inflammasomes, but exactly how CTB produces IL-1β remains unclear.

CTB, which binds to ganglioside GM1 on the cell surface, is incorporated within the cell and translocated from Golgi apparatus to the endoplasmic reticulum (ER) through the retrograde transport pathway (*10–13*). CTB is retained in ER and induces ER stress responses. However, previous studies have been performed using in *in vitro* cell lines. Whether or how ER stress responses are involved in IL-1β production, especially in *in vivo* macrophages, has been unclear.

In this study, we analyzed effects of CT or CTB on RPMs. In RPMs, incorporated CTB was retained in ER and induced expression of ER-stress related genes as well as IL-1β production in a GM1-dependent manner. Pharmacological inhibition of an ER-stress sensor IRE1α and genetic deletion of *Ern1* encoding IRE1α caused decrease of IL-1β production from RPMs in response to CT or CTB. Our study first showed the critical roles of IRE1α in bacterial exotoxin-induced inflammasome activation in tissue-resident macrophages.

## RESULTS

### CT induces the expression of ER stress-related genes in a ganglioside GM1-dependent manner from LPS-primed RPMs

We first examined whether CT, CTA or CTB can activate RPMs to produce IL-1β. CTA could not induce IL-1β production from RPMs even in the presence of LPS. CT as well as CTB could induce IL-1β production when RPMs were prestimulated with LPS O55:B5 or LPS O111:B4 (fig. S1A). Similar to CTB, CT could induce IL-1β production from RPMs in synergy with LPS O55:B5 in a manner dependent on GM1, NLRP3 or pyrin (fig. S1, B to E). Thus, both CT and CTB can induce IL-1β production from LPS-primed RPMs through NLRP3 and pyrin inflammasome. Hereafter, LPS O55:B5 is used when referring to LPS, unless otherwise stated.

We performed comprehensive analysis of CT- or CTB-induced genes in LPS-primed RPMs (Fig. 1). In response to CT, expression of 828 genes was upregulated more than 2-fold (Fig. 1A). Enrichment analysis of these genes showed activation of 14 pathways, among which the “protein processing in endoplasmic reticulum” pathway was the most enriched pathway (*P*=1.1E-20) (Fig. 1B). The pathway is comprised of 42 ER stress-related genes, which contain 24 X-box binding protein 1 (XBP1) target genes (*14–16*). In response to CTB, expression of 510 genes was upregulated more than 2-fold and 15 pathways were enriched by pathway analysis. Similar to the responses to CT, the “protein processing in endoplasmic reticulum” pathway was the most enriched pathway (*P*=8.6E-8) (Fig. 1, C and D).

**Fig. 1.**
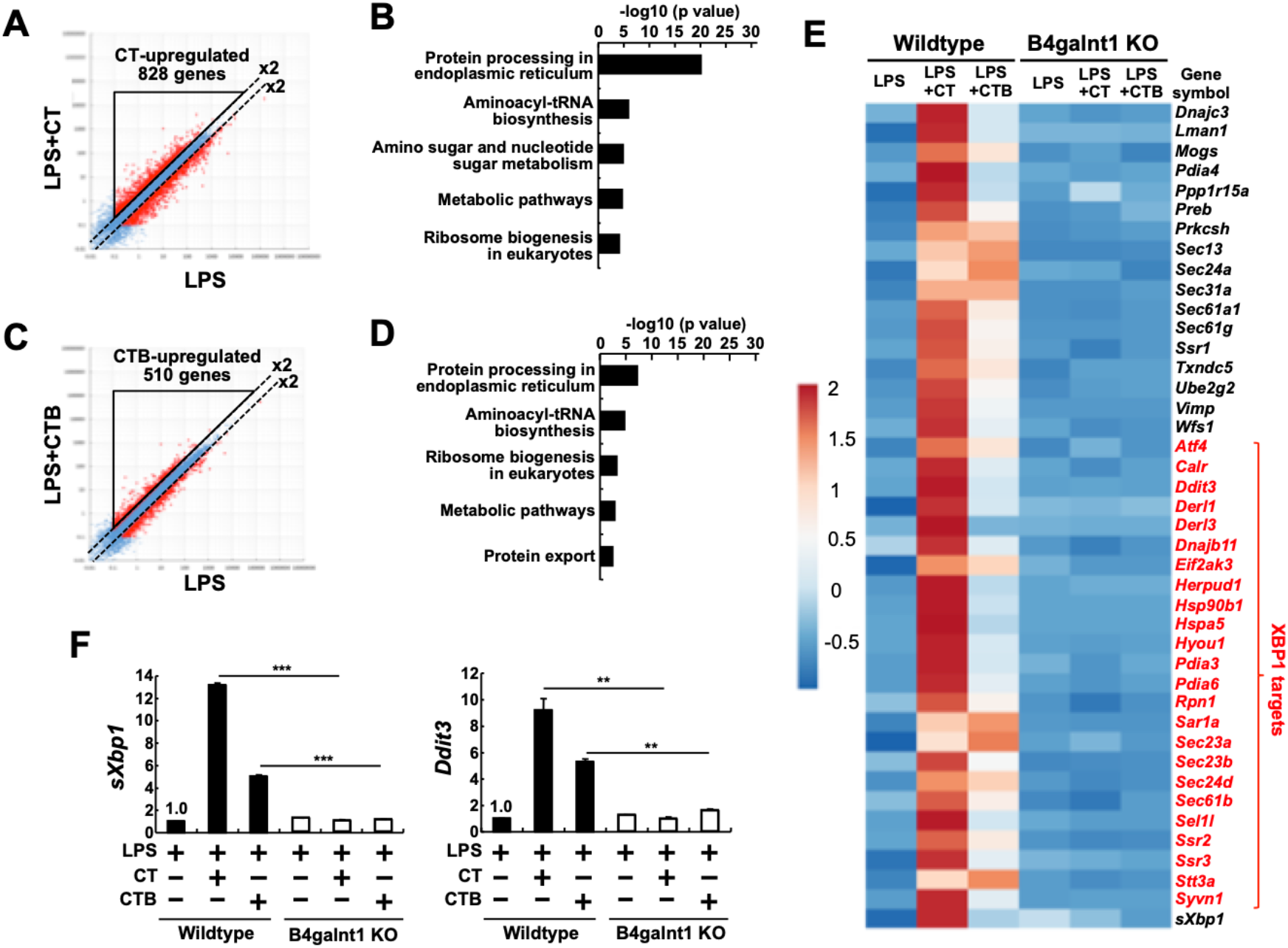
CT induces the ER stress responses in LPS-primed RPMs through a GM1-dependent manner. **(A-F)** RPMs from littermate B4galnt1 wild-type mice or B4galnt1-deficient mice were first cultured for 5 h in the presence of 500 ng ml^−1^ LPS O55:B5. Then 20 µg ml^−1^ CT or CTB were added and further cultured for 19 h. Total RNAs were subjected to RNAseq **(A-E)** or quantitative real-time PCR **(F)**. **(A)** Scatter plot analysis of LPS or LPS+CT-stimulated wild-type RPMs by RNAseq. The lines indicate the 2-fold difference. **(B)** KEGG pathway analysis of CT-upregulated genes in **(A)**. **(C)** Scatter plot analysis of LPS or LPS+CTB-stimulated wild-type RPMs by RNAseq. Lines indicate the 2-fold difference. **(D)** KEGG pathway analysis of CTB-upregulated genes in **(C)**. **(E)** Heatmap of ER-stress related genes in response to indicated stimuli in wild-type or B4galnt1-deficient RPMs. **(F)** The expression of representative ER-stress related genes. RNAs were subjected to quantitative real-time PCR analysis for *sXbp1* and *Ddit3*. Data are representative of three independent experiments. The results are presented as means ± SD. \*\**P* < 0.01, \*\*\**P* < 0.001.

ER stress is triggered by accumulation of unfolded proteins in the ER lumen. These proteins are recognized by ER stress sensors, which contain inositol-requiring enzyme 1-alpha (IRE1α), Protein kinase R (PKR)-like endoplasmic reticulum kinase (PERK) and ATF6. These sensors induce various kinds of ER-stress-related genes and mediate ER stress responses or unfolded protein responses (UPRs). In GM1-deficient RPMs, expression of 621 and 425 genes was upregulated more than 2-fold in responses to CT and CTB, respectively, but none of 42 ER stress-related genes were upregulated by CT or CTB (Fig. 1E). Quantitative RT-PCR analysis verified that CT or CTB can induce spliced form of *Xbp1* mRNA (*sXbp1*) and expression of *Ddit3*, which are downstream of IRE1α and PERK signaling, respectively (Fig. 1F). These results suggest that CT as well as CTB can activate ER stress sensors in a GM1-dependent manner from LPS-primed RPMs.

### CT and Tunicamycin (Tm) induces expression of similar sets of ER stress-related genes in resting RPMs

The effects of CT alone were then compared with those of an ER stress inducer, tunicamycin (Tm). Expression of 677 and 941 genes was upregulated more than 2-fold in CT- and Tm-stimulated RPMs (Fig. 2, A and C). According to the enrichment analysis, Tm induced activation of 16 pathways, among which the “protein processing in endoplasmic reticulum” pathway was the most enriched pathway (*P*=3.9E-32 and *P*=2.4E-24 for CT and Tm stimulation, respectively) (Fig. 2, B and D). Comparison of CT-stimulated and Tm-stimulated genes revealed that 423 genes containing XBP1 target genes were overlapped (Fig. 2E). Pathway analyses were then performed for these overlapped genes and the “protein processing in endoplasmic reticulum” pathway was most prominently enriched (*P*=8.6E-31 Fig. 2, F to H). Quantitative RT-PCR analysis verified that expression of both *sXbp1* and *Ddit3* mRNAs was induced by CT or Tm, but not LPS (Fig. 2I). These results indicate that CT can activate RPMs to induce expression of ER-stress related genes in similar to Tm. Next, we investigated whether CT or Tm induces the expression of XBP1s, encoded by *sXbp1* in RPMs (Fig. 2, J and K). CT or Tm upregulated XBP1s expression in wild-type RPMs (Fig. 2, J and K). In GM1-deficient RPMs, CT-induced XBP1s upregulation was completely abolished but the Tm-induced XBP1s upregulation was intact (Fig. 2, J and K). These results suggest that CT induces IRE1α signaling leading to *Xbp1* splicing in a GM1-dependent manner.

**Fig. 2.**
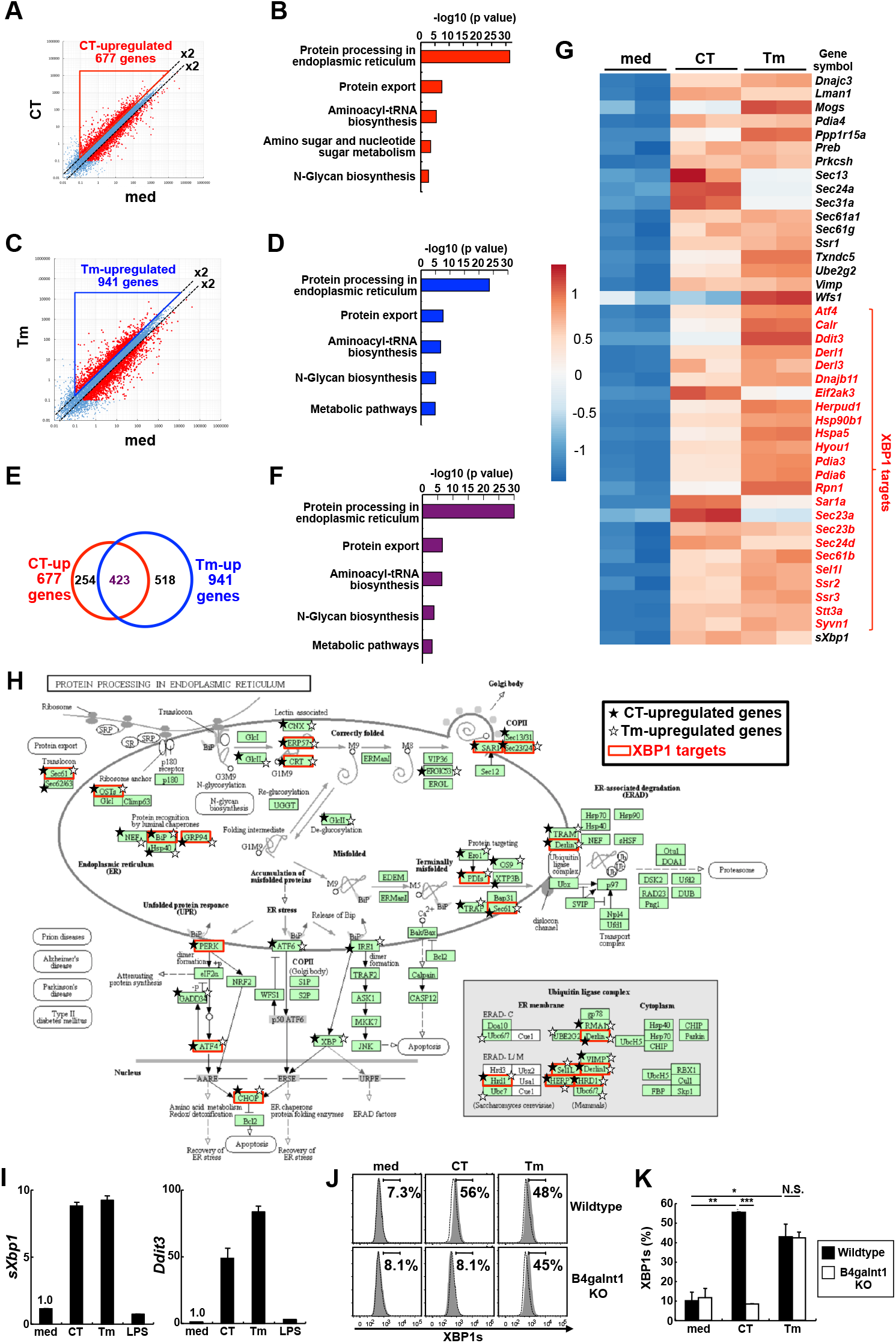
CT as well as Tm induces the expression of similar sets of ER stress-related genes. **(A-I)** RPMs from wild-type C57BL/6 mice were first cultured for 5 h in the presence or absence of 500 ng ml^−1^ LPS O55:B5. Then, 20 µg ml^−1^ CT, 20 µg ml^−1^ CTB or 10 µg ml^−1^ Tm were added and further cultured for 19 h. Total RNAs were subjected to RNAseq **(A-H)** or quantitative real-time PCR **(I)**. **(A)** Scatter plot analysis of non-stimulated or CT-stimulated RPMs by RNAseq. Lines indicate the 2-fold difference. **(B)** KEGG pathway analysis of CT-upregulated genes in (A). **(C)** Scatter plot analysis of non-stimulated or Tm-stimulated RPMs by RNAseq. The lines indicate the 2-fold difference. **(D)** KEGG pathway analysis of Tm-upregulated genes in **(C)**. **(E)** Venn diagram depicts the numbers of genes upregulated by CT and Tm in RPMs compared to non-stimuli. **(F)** KEGG pathway analysis of overlapped genes in (E). **(G)** Heatmap of ER-stress related genes in response to indicated stimuli in RPMs (N=2). **(H)** KEGG pathway visualization of Protein processing in endoplasmic reticulum associated with CT and Tm-upregulated genes. Black stars and white stars show CT-upregulated genes and Tm-upregulated genes, respectively. Genes, which were coded red, represent XBP1 targets. **(I)** Expression of representative ER-stress related genes. RNAs were subjected to quantitative real-time PCR analysis for *sXbp1* and *Ddit3*. Data are representative of two independent experiments. The results are presented as means ± SD. **(J, K)** rPECs from littermate B4galnt1 wild-type mice or B4galnt1-deficient mice were cultured with or without 20 µg ml^−1^ CT or 10 µg ml^−1^ Tm for 19 h. Cells were subjected to flow cytometric analysis for detection of XBP1s expression. **(J)** Histograms of XBP1s from wild-type or B4galnt1-deficient LIVE/DEAD^−^ F4/80^+^ CD11b^+^ rPECs in response to indicated stimuli are shown. Cells were harvested and subjected to intracellular staining of transcription factor XBP1s. Shaded histograms and dotted lines represent the staining for anti-XBP1s PE and control isotype mouse IgG1 PE antibodies (Cat:400112, Biolegend), respectively. Numbers indicate the percentages of gated cells among LIVE/DEAD^−^ F4/80^+^ CD11b^+^ rPECs. Data are representative of two independent experiments. **(K)** Percentages of XBP1s expression in (J) (N=2 mice/ genotype). The results are presented as means ± SD. \**P* < 0.05, \*\**P* < 0.01, \*\*\**P* < 0.001. N.S. represent not significant. Data are representative of two independent experiments.

### CT is incorporated into ER in a GM1-dependent manner

Next, we investigated the intracellular localization of incorporated CT subunits: CTA, CTB and CTA1, which is a proteolytic cleavage product of CTA. They were detected mainly in ER, rather than in the cytosol (Fig. 3A). In GM1-deficient macrophages, neither CTA, CTB nor CTA1 were present in ER or cytosol (Fig. 3A). We have also analyzed intracellular localization of CTB by fluorescence microscopy. IRE1α, an ER stress sensor, was co-stained with an ER marker DiOC6, consistent with its localization in ER (fig. S2). In wild-type RPMs, FITC-labeled CTB was colocalized with IRE1α, but the co-localization was abolished in GM1-deficient RPMs (Fig. 3, B and C). These results suggest that CTB is incorporated into ER in a GM1-dependent manner in RPMs.

**Fig. 3.**
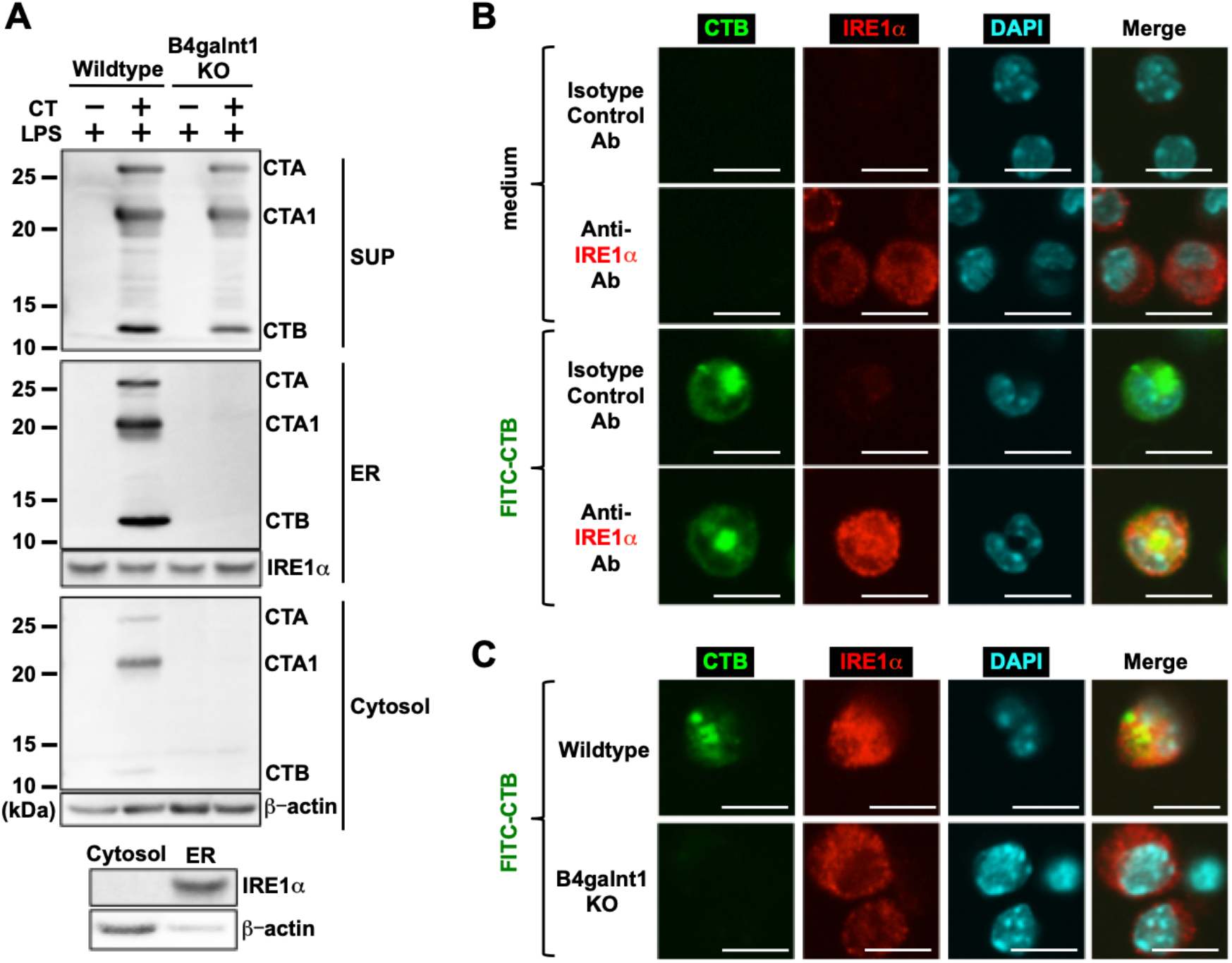
CT is translocated to the ER in a GM1-dependent manner. **(A)** rPECs from littermate B4galnt1 wild-type mice or B4galnt1-deficient mice were first cultured for 5 h in the presence of 500ng ml^−1^ LPS O55:B5. Then 20 µg ml^−1^ CT were added and further cultured for 6 h. Cell fractions were then subjected to western blot analysis for antibodies against CT, IRE1α or β-actin. IRE1α and β-actin were used as the ER and cytosol markers, respectively. **(B)** RPMs from wild-type C57BL/6 mice were treated with or without 1 µg ml^−1^ of FITC-conjugated CTB for 12 h. Cells were then subjected to confocal microscopic analysis. Green, Red and Blue coloring represent CTB internalization, IRE1α and DAPI staining, respectively. Rabbit IgG was used as Isotype control Ab. Scale bars represent 10µm. Data are representative of two independent experiments. **(C)** RPMs from littermate B4galnt1 wild-type mice or B4galnt1-deficient mice were treated with 1 µg ml^−1^ of FITC-conjugated CTB for 12 h. Cells were then subjected to confocal microscopic analysis as described in **(B)**. Data are representative of two independent experiments.

### CT induces *Xbp1* mRNA splicing through IRE1α

We analyzed how ER stress sensors are involved in CT-induced the expression of ER stress-related genes in RPMs (Fig. 4). We first utilized 4µ8c, an IRE1α inhibitor, which inhibits both kinase and RNase activities of IRE1α, by interacting with the kinase active site residue, Lys599, in IRE1α (*17*). In the presence of 4µ8c, CT- or CTB-induced *Xbp1* mRNA splicing in LPS-primed RPMs was severely impaired (Fig. 4A). We have also examined the effects of GSK2850163, another IRE1α inhibitor, which also inhibits both kinase and RNase activities of IRE1α through the binding to Lys599 in IRE1α (*18*). GSK2850163 inhibited CT- or CTB-induced *Xbp1* splicing. Meanwhile, GSK2606414, a PERK inhibitor, (*19*) inhibited *Ddit3* expression induced by CT or CTB (Fig. 4A). CT or CTB can therefore activate both IRE1α and PERK signaling.

**Fig. 4.**
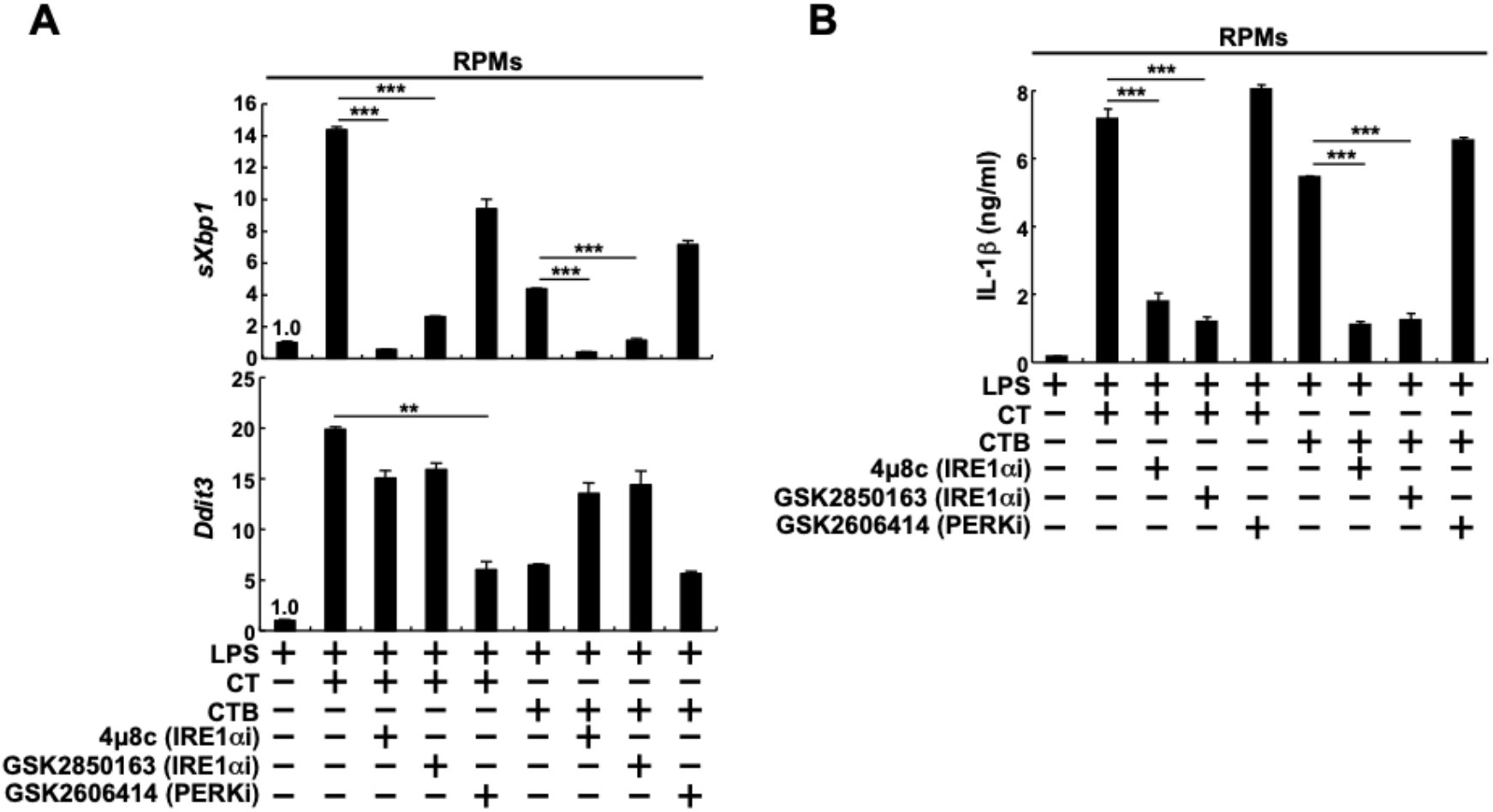
CT induces *Xbp1* splicing in LPS-primed RPMs through IRE1α. **(A,B)** RPMs from wild-type C57BL/6 mice were first cultured for 5 h in the presence of 500 ng ml^−1^ LPS O55:B5. Then 20 µg ml^−1^ CT or CTB were added and further cultured for 19 h. IRE1α inhibitor, 4µ8c (36 µM) and GSK2850163 (40 µM), PERK inhibitor GSK2606414 (3 µM), or DMSO as control were added 3h before addition of CT or CTB. Total RNAs were subjected to quantitative real-time PCR analysis for *sXbp1* and *Ddit3* **(A)**. Culture supernatants were subjected to ELISA for IL-1β **(B)**. Results are presented as means ± SD. \*\**P* < 0.01, \*\*\**P* < 0.001.

We then analyzed whether IRE1α and/or PERK are involved in CT-induced IL-1β production from LPS-primed RPMs (Fig.4B). In the presence of an IRE1α inhibitor 4µ8c or GSK2850163, CT- or CTB-induced IL-1β production was decreased in LPS-primed RPMs (Fig. 4B). Meanwhile, GSK2606414, a PERK inhibitor, did not inhibit IL-1β production induced by CT or CTB (Fig. 4B). These results indicate that IRE1α, but not PERK, is involved in CT or CTB-induced IL-1β production from LPS-primed RPMs.

### IRE1α is required for CT or CTB-induced IL-1β production from LPS-primed RPMs

To further investigate the involvement of IRE1α in CT- or CTB-induced effects, we examined macrophage-specific IRE1α-deficient (Mac-IRE1α KO) mice. In Mac-IRE1α KO mice, the IRE1α-RNase domain, which is involved in *Xbp1* splicing, is deleted by cre recombinase, expression of which is driven by *Lyz2* promoter (*20, 21*). Littermates expressing cre recombinase and carrying one floxed *Ern1* locus were used as control. We verified that this deletion was detected at protein levels in RPMs, but not in non-RPMs (fig. S3). In Mac-IRE1α KO mice, *sXbp1* induction was significantly decreased in RPMs, although it was retained in non-RPMs (Fig. 5A). Meanwhile, CT- or CTB-induced *Ddit3* upregulation, which is mediated by PERK signaling, was retained in RPMs from Mac-IRE1α KO mice (Fig. 5A). We then investigated expression of XBP1s, encoded by *sXbp1* in RPMs (Fig. 5, B and C). In Mac-IRE1α KO RPMs, both CT- and Tm-induced XBP1s upregulation was decreased compared with that in control RPMs (Fig. 5, B and C). These results indicate that IRE1α is required for CT- or Tm-induced *Xbp1* splicing and XBP1s expression.

**Fig. 5.**
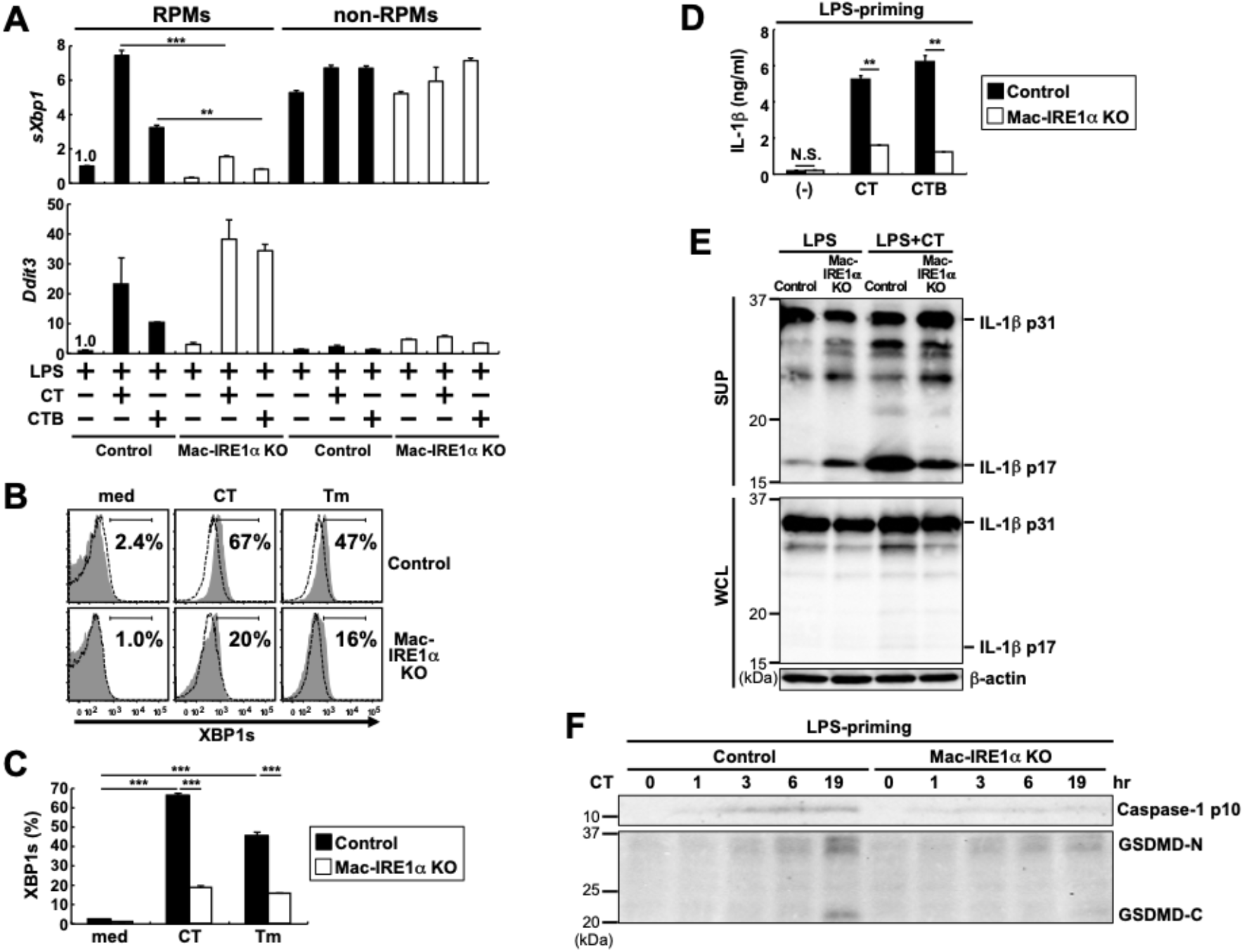
IRE1α is required for CT or CTB-induced IL-1β production from LPS-primed RPMs. **(A)** RPMs from littermate *Ern1^+/fl^*, *Lyz2*^+/cre^ mice or *Ern1^fl/fl^*, *Lyz2*^+/cre^ mice were first cultured for 5 h in the presence of 500 ng ml^−1^ LPS O55:B5. Then 20 µg ml^−1^ CT or CTB were added and further cultured for 19 h. Littermate *Ern1^+/fl^*, *Lyz2*^+/cre^ mice and *Ern1^fl/fl^*, *Lyz2*^+/cre^ mice were used as control and Mac-IRE1α KO mice, respectively. Total RNAs were subjected to quantitative real-time PCR analysis for *sXbp1* and *Ddit3*. **(B,C)** rPECs from control or Mac-IRE1α KO mice were cultured with or without 20 µg ml^−1^ CT or 10 µg ml^−1^ Tm for 19 h. Cells were subjected to flow cytometric analysis for detection of XBP1s expression as described in the Figure 2J legend. **(B)** Histograms of XBP1s from control or Mac-IRE1α KO LIVE/DEAD^−^ F4/80^+^ CD11b^+^ rPECs in response to indicated stimuli are shown. Data are representative of two independent experiments. **(C)** Percentage of XBP1s expression in **(B)** (N=2 mice/ genotype). Results are presented as means ± SD. \*\*\**P* < 0.001. Data are representative of two independent experiments. **(D)** RPMs from control or Mac-IRE1α KO mice were cultured as described in **(A)** legend. Culture supernatants were subjected to ELISA for IL-1β. Results are presented as means ± SD. \*\**P* < 0.01. N.S. = not significant. **(E)** RPMs from control or Mac-IRE1α KO mice were first cultured for 5 h in the presence of 500 ng ml^−1^ LPS O55:B5. Then, 20 µg ml^−1^ CT was added and further cultured for 19 h. WCL and culture supernatants (SUP) were subjected to western blot analysis with anti-IL-1β or anti-β actin-antibodies. **(F)** RPMs from control or Mac-IRE1α KO mice were first cultured for 5 h in the presence of 500 ng ml^−1^ LPS O55:B5. Then, 20 µg ml^−1^ CT were added and cultured. Culture supernatants were harvested at the indicated time points and subjected to western blot analysis with anti-Caspase-1 p10 or GSDMD antibodies. Data are representative of two independent experiments.

We then investigated CT- or CTB-induced IL-1β production in RPMs. CT- or CTB-induced IL-1β production was significantly impaired in LPS-primed Mac-IRE1α KO RPMs (Fig. 5D). Western blot analysis showed decrease of mature IL-1β protein (p17) and relative increase of pro-IL-1β protein (p31) in CT-stimulated IRE1α KO RPMs, compared with CT-stimulated control RPMs (Fig. 5E). These results suggest that IRE1α is required for CT-induced inflammasome activation leading to IL-1β processing.

Inflammasome activation induces caspase cascades leading to cleavage of Gasdermin D (GSDMD) to GSDMD N-terminal fragment (GSDMD-N), which generates membrane pore formation and induces IL-1β secretion. We then investigated whether IRE1α is involved in this step. In LPS-primed control RPMs, both the p10 subunit of active caspase-1 (Caspase-1 p10) and GSDMD-N were induced at 3 hr after CT stimulation and further increased until 19 hr. Meanwhile, in CT-stimulated LPS-primed Mac-IRE1α KO RPMs, induction of both Caspase-1 p10 and GSDMD-N was impaired throughout the observed periods (Fig. 5F). Thus, these data suggest that IRE1α is required for CT-induced caspase-1 activation and GSDMD processing.

### IRE1α is required for activation of both NLRP3 and pyrin inflammasomes

CT or CTB can activate both NLRP3 and pyrin inflammasomes. It, however, is unclear which inflammasome is activated through IRE1α. Adenosine triphosphate (ATP) and *Clostridium difficle* toxin B (TcdB) could activate LPS-primed RPMs to produce IL-1β in NLRP3- and pyrin-dependent manner, respectively (8). Compared with control RPMs, Mac-IRE1α KO RPMs showed impaired responses to both ATP, and TcdB (Fig. 6, A and B), indicating that IRE1α was required for both NLRP3 and pyrin inflammasome activation.

**Fig. 6.**
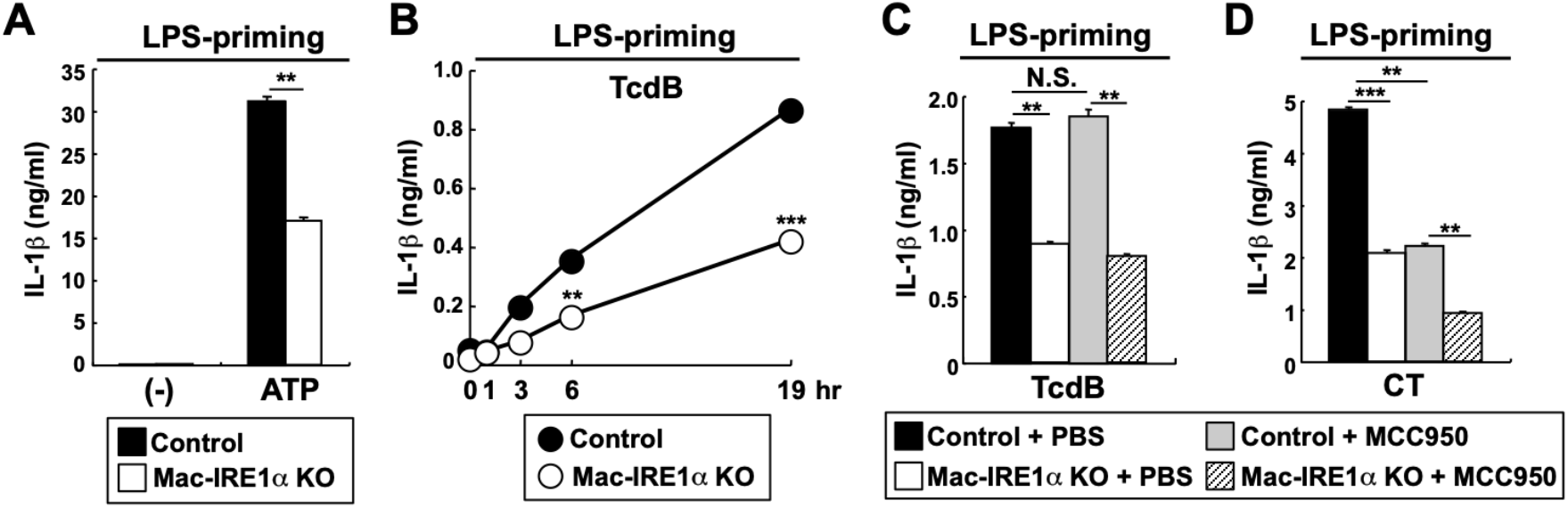
IRE1α is required for NLRP3 or pyrin-mediated IL-1β production from LPS-primed RPMs. **(A-D)** RPMs from control or Mac-IRE1α KO mice were first cultured for 5 h in the presence of 500 ng ml^−1^ LPS O55:B5. **(A)** 1 mM ATP was added and further cultured for 19 h. **(B)** 5 µg ml^−1^ TcdB were added and cultured. Culture supernatants were harvested at the indicated time points. **(C,D)** 5 µg ml^−1^ TcdB **(C)** or 20 µg ml^−1^ CT **(D)** was added and cultured for 19 h. NLRP3 inhibitor, MCC950 (1 µM) or PBS as vehicle control were added 3 h before addition of TcdB, CT. IL-1β production was measured by ELISA. The bars or symbols represent means ± SD. \*\**P* < 0.01, \*\*\**P* < 0.001. N.S. = not significant.

We further tested whether CT-induced IL-1β production through pyrin inflammasome involves IRE1α in the presence of NLRP3-specific inhibitor. ATP-, but not TcdB-induced IL-1β production was abolished by MCC950 (fig. S4 and Fig. 6C), verifying that MCC950 could inhibit NLRP3, but not pyrin, inflammasome. In the presence of MCC950, Mac-IRE1α KO RPMs showed less production of IL-1β than control RPMs, in response to CT as well as TcdB (Fig. 6, C and D). The results indicate that IRE1α is required for CT-induced activation of pyrin inflammasome.

## DISCUSSION

We clarified the role of IRE1α in CT-induced effects in murine macrophages (Fig.7). CT, incorporated with GM1, is translocated to the ER and activates IRE1α, an ER stress sensor which can lead to *Xbp1* mRNA splicing. Inhibitors and knockout mice analysis indicated that IRE1α is required for CT-induced IL-1β production from resident peritoneal macrophages.

**Fig. 7.**
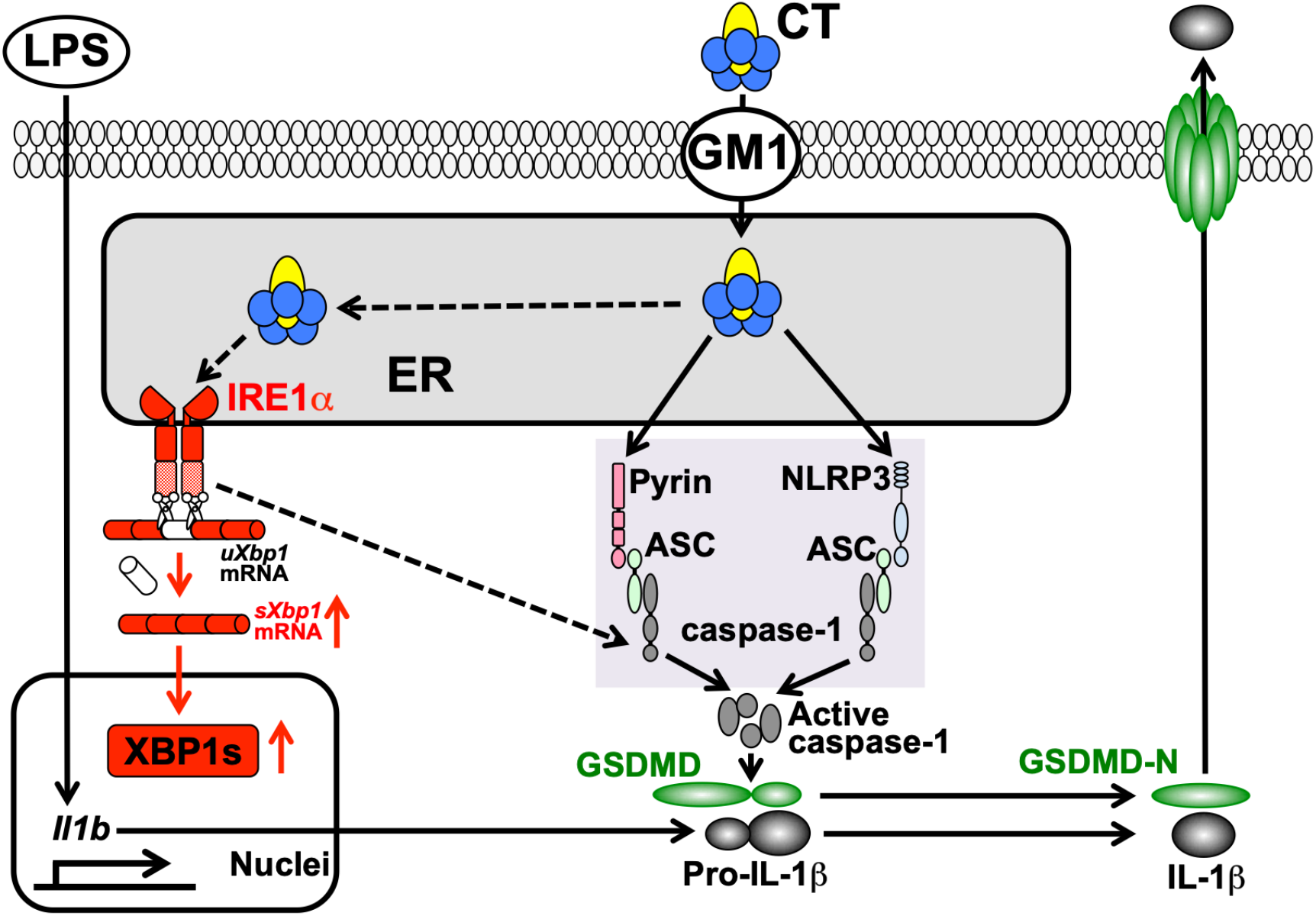
A hypothetical model of CT-induced IL-1β production in resident peritoneal macrophages. CT, incorporated with GM1, translocates to the ER and activates an ER stress sensor IRE1α. Then, IRE1α is required for Pyrin or NLRP3-mediated IL-1β production through caspase-1 activation and GSDMD processing.

CT or CTB can activate both NLRP3 and pyrin inflammasomes (*8*)(fig. S1). IRE1α inhibitors attenuate ER-stress activator-induced NLRP3 inflammasome activation in a human monocytic leukemia cell line or murine myeloid cells generated *in vitro* (*22, 23*). Furthermore, analysis on IRE1α KO mice also show that a saturated fatty acid, which functions not only as an ER-stress activator but also as an NLRP3 inflammasome activator, induces IL-1β production from LPS-prestimulated murine BM-derived GM-CSF-induced myeloid cells in an IRE1α-dependent manner (*15*). Consistent with these studies, our findings clarify that CT, which can induce ER-stress responses, produces IL-1β in an IRE1α dependent manner in murine peritoneal macrophages (Fig.5, D to F).

Pyrin inflammasome is activated by a variety of toxins. They modify the RhoA GTPase, which regulates cytoskeletal remodeling, either through adenylylation, glucosylation, deamidation or ADP-ribosylation and inactivate it, thereby leading to pyrin inflammasome activation (*24–26*). In response to TcdB, a pyrin inflammasome activator, IRE1α KO macrophages produced less IL-1β than control macrophages (Fig.6, B and C), indicating that IRE1α is required for pyrin inflammasome activation. TcdB enters cells through endocytosis, moves to the acidic endosomal compartment, and then releases the N-terminal glucosyltransferase domain (GTD) by autoproteolysis into the cytosol (*27–29*). GTD then moves to the plasma membrane, where it inactivates RhoA by glucosylation of Thr37 in RhoA (*25, 26*). It remains unclear at present whether TcdB translocates to the ER or induces the ER stress. Our findings, however, are the first to demonstrate that IRE1α is required for pyrin inflammasome activation.

It can be assumed that IRE1α targets a common pathway between NLRP3 and pyrin inflammasome activation. Consistently, both caspase-1 activation and GSDMD processing, which are required for IL-1β production in response to NLRP3 and pyrin inflammasome activation (*30–33*), were impaired in the absence of IRE1α (Fig. 5F). While the pathways upstream of NLRP3 and pyrin inflammasomes do not overlap, the idea that IRE1α signaling is linked to the upstream pathways of each inflammasome cannot be excluded. Further study is required to clarify how IRE1α is linked to inflammasome activation, but this study clearly demonstrates that IRE1α signaling is required for the NLRP3 and pyrin inflammasome activation in *in vivo* macrophages. Our present results here demonstrate the critical roles of IRE1α in CT- or CTB-induced IL-1β production in murine resident macrophages. The findings should contribute not only to the understanding of the molecular mechanisms on CT-induced immune adjuvant effects but also to clarification of novel roles of IRE1α in immune responses and pathogenesis of autoinflammatory disease involved in NLRP3 or pyrin inflammasome activation.

## MATERIALS AND METHODS

### Study design

This study aimed to clarify a mechanism of CT-induced IL-1β production from LPS-primed murine resident peritoneal macrophages. We performed RNA-seq and found that enhancement of unfolded protein responses in CT-stimulated RPMs in a GM1-dependent manner. We used Mac-IRE1α KO mice and UPRs sensor inhibitors to investigate the role of UPRs in CT-stimulated RPMs. Experiments were conducted in replicates as indicated in the Figure Legends. Age-matched and sex-matched mice were used for all experiments. Experiments were not conducted in a blinded or randomized manner, and no outliers were excluded.

### Reagent

LPS O55:B5 (L2637) and LPS O111:B4 (L3012) were purchased from Sigma-Aldrich. CTB (Choleragenoid, 103B) and CT (100B) were purchased from List Biological Laboratories. ATP (tlrl-atp) was purchased from InvivoGen. TcdB (6246-GT-020) was purchased from R&D systems. Tunicamycin (654380-10MGCN) was purchased from Calbiochem (Merck).

### Mice

Eight-week to 18-week old C57BL/6 mice were purchased from CLEA Japan. β1,4-*N*-acetylgalactosaminyltransferase (B4galnt1)-deficient mice, which lack all complex gangliosides including GM1, have been described previously, they were backcrossed more than seven times with C57BL/6 mice (*34, 35*). An adaptor, apoptosis-associated speck-like protein containing CARD (ASC)-deficient and NLRP3-deficient mice have been described previously (*36–38*). Pyrin-deficient mice have been described previously (*8*). Floxed IRE1α (*Ern1^fl/fl^*) mice have been described previously (*20*) and crossed with the Lysozyme 2-Cre (*Lyz2*^+/cre^) line (*21*) (JACKSON B6 129P2-Lyz2^tm1(cre)Ifo^ /J Stock No.004781 The Jackson Laboratory JAX: 004781) to generate macrophage-specific IRE1α deficient mice (*Ern1^fl/fl^*, *Lyz2*^+/cre^) as Mac-IRE1α KO mice. Then, littermate *Ern1^+/fl^*, *Lyz2*^+/cre^ mice were used as control mice. All mice were bred and maintained in the Animal Facility of Wakayama Medical University or Osaka University under specific pathogen-free conditions and were used according to the institutional guidelines of Wakayama Medical University and Osaka University. All animal experiments were approved by the animal research committees and were carried out in accordance with approved guidelines of the animal care committees of Wakayama Medical University and Osaka University.

### Cell preparation

Cells were harvested from a peritoneal cavity with PBS and used as resident peritoneal exudate cells (rPECs). RPMs (F4/80^+^ cells) or non-RPMs (F4/80^−^ cells) were isolated from rPECs by using magnetic-activated cell sorting (MACS). The details were described in our previous paper (*8*). Briefly, rPECs were blocked with anti-CD16/32 (2.4G2), followed by the incubation with biotinylated anti-F4/80 (Cl:A3-1,Serotec). After washing cells were further incubated with streptavidin micro-beads (Miltenyi Biotec) and then magnetically sorted with LS columns as per the manufacture’s instructions (Miltenyi Biotec). The flow through cells were collected and used as non-RPMs (F4/80^−^ cells). Micro-beads conjugated cells were collected and used as RPMs (F4/80^+^ cells). Purity of RPMs was typically 85-90%.

### Measurement of IL-1β production

For measuring IL-1β production *in vitro*, 8 × 10^4^ cells per well in 96-well plates were first cultured for 5h in the absence or presence of LPS O55:B5 or LPS O111:B4. CT, CTB, ATP or TcdB were then added at indicated concentrations and further cultured for indicated 19h. IRE1α inhibitor, 4µ8c (14003-96-4, Cayman), GSK2850163 (SML-1684-5MG, Sigma-Aldrich), PERK inhibitor GSK2606414 (5107, Tocris) and NLRP3 inhibitor MCC950 (S7809, Selleck) were added 3h before addition of CT, CTB. Dimethyl sulfoxide (DMSO) (13408-64, Nacalai Tesque) was used as a solvent control for inhibitors. Culture supernatants were subjected to enzyme-linked immunosorbent assay (ELISA) for IL-1β (SMLB00C, R&D systems). The assays were performed as recommended by the manufacturers.

### RNA sequencing

RPMs (4 × 10^5^ cells) were harvested and total RNA was extracted with RNeasy micro kit (QIAGEN). RNA libraries were prepared using TruSeq stranded mRNA Library Prep Kit (Illumina). Sequencing was performed on HiSeq 2500 platform in a 75-base single-end mode or NovaSeq 6000 platform in a 101-base single-end mode. Illumina RTA 1.18.64 software (for HiSeq 2500) or Illumina RTA 3.4.4 software (for NovaSeq 6000) was used for base calling. Generated reads were mapped to the mouse (mm10) reference genome using TopHat v2.1.1 in combination with Bowtie2 ver. 2.2.8 and SAMtools ver. 0.1.18. Fragments per kilobase of exon per million mapped fragments (FPKMs) were calculated using Cuffdiff 2.2.1. The Subio Platform and Subio Basic Plug-in (version 1.20; Subio) were used to depict scatter plots.

The DAVID Bioinfomatics database (*39*) and Kyoto Encyclopedia of Genes and Genomes (KEGG) (*40*) annotations were used for enrichment analysis. The web tool ClustVis (*41*) was used to draw heatmap.

### Quantitative real-time PCR

Using PrimeScript RT reagent Kit (TaKaRa), 0.5 µg of total RNAs were reverse transcribed into complementary DNA. Relative expression levels of RNA transcripts were determined using gene-specific primers, TB Green Premix Ex Taq II (TaKaRa) and StepOneplus Real-Time PCR system (Applied Biosystems). TaqMan probes (TaqMan Gene Expression Assay,Applied Biosystems) were used for 18S rRNA (internal control). The expression of all genes was normalized to that of 18S rRNA and is represented as the ratio to the indicated reference samples. Gene specific primers used in this study were as follows: *sXbp1* (*42*), sense, 5’-AAGAACACGCTTGGGAATGG-3’, antisense, 5’-CTGCACCTGCTGCGGAC-3’; *Ddit3* (*43*), sense, 5’-CCCAGGAAACGAAGAGGAAG-3’, antisense, 5’-AGTGCAGTGCAGGGTCACAT-3’. All primers were validated for linear amplification.

### Western blot analysis

Five hundred thousands cells of RPMs were lysed in radioimmunoprecipitation assay (RIPA) buffer containing 50 mM Tris-HCl at pH 8.0, 150 mM NaCl, 1 % NP-40, 0.5 % sodium deoxycholate, 0.1 % sodium dodecyl sulfate (SDS) with protease inhibitor cocktail (11836153001, Roche). Whole cell lysate (WCL) samples or culture supernatants were then boiled with SDS sample Buffer and subjected to western blot analysis.

ER and cytosol fractions from rPECs were prepared using the Mem-PER Plus Membrane Protein Extraction Kit (89842, Thermo Fisher Scientific). Briefly, cells were harvested and resuspended with permeabilization buffer containing protease inhibitor cocktail (11836153001, Roche) and incubated for 10 min on ice with constant mixing. After centrifugation at 16,000 *g* for 15 min, supernatants were collected as Cytosol fractions. The pellets were resuspended with 100 μl of solubilization buffer containing protease inhibitor cocktail (11836153001, Roche) and incubated for 30 min on ice with constant mixing. After centrifugation at 16,000 *g* for 15 min, supernatants were collected as ER fractions. The isolated ER and Cytosol fractions were boiled in the SDS sample buffer and subjected to Western blot analysis.

Protein samples were separated in polyacrylamide gel and then transferred to polyvinylidene difluoride membranes (1620177, Bio-Rad) or nitrocellulose membrane (1620146, Bio-Rad) for detecting caspase-1 p10. These membranes were incubated with primary antibodies, followed by secondary antibodies conjugated to horseradish peroxidase (HRP).

The following antibodies (Abs) were used: anti-β-actin Ab (C4) (sc-47778; 1/1000 dilution, Santa Cruz Biotechnology), anti-IRE1α (3294; 1/1000 dilution, cell signaling technology), anti-CT (ab123129; 1/1000 dilution, abcam), biotinylated anti-IL-1β (BAF401; 1/250 dilution: R&D Systems), anti-Caspase-1 p10 (sc-514; 1/1000 dilution, Santa Cruz Biotechnology), anti-GSDMD (ab209845; 1/1000 dilution, abcam), anti-mouse IgG-HRP (NA931V; 1/1000 dilution, GE Healthcare), anti-rabbit IgG-HRP (NA934V; 1/1000 dilution, GE Healthcare) and Streptavidin-HRP Conjugate (RPN1231; 1/250, Merck). Western Lightning Plus-ECL (PerkinElmer) was used for positive signals and the chemiluminescence was detected using a Light Capture AE-6971/2 device (ATTO) or ChemiDoc Touch Imaging System (Bio-Rad).

### Flow cytometry

rPECs were incubated with CT (20 µg ml^−1^) or Tm (10 µg ml^−1^) for 19h. Cells were then harvested, incubated with anti-CD16/32 (clone 2.4G2, cat:70-0161-M001, TONBO biosciences) to block Fc receptors and stained with biotinylated anti-mouse F4/80 (clone: Cl:A3-1, cat:MCA497B, BIO-RAD), PerCP-Cy5.5-conjugated streptavidin (551419, BD Biosciences) and V450-conjugated anti-CD11b (clone:M1/70, cat:560456, BD Biosciences). For intracellular staining of transcription factor XBP1s, cells were fixed and permeabilized with Foxp3/Transcription Factor Staining Buffer Set (00-5523, Invitrogen), then stained with anti-XBP-1S-PE (clone: Q3-695, cat:562642, BD Biosciences) or control isotype mouse IgG1 PE (400112, Biolegend) antibodies. Dead cells, which can be detected as positive cells by AmCyan channel, were excluded using the LIVE/DEAD Fixable Dead Cell Aqua Stain Kit (L34966, Invitrogen). Stained cells were analyzed on a FACS Verse and data were processed with FlowJo Version 8.8.7 software (TreeStar).

### Subcellular localization of CT

CELLview cell culture dish (627975, Greiner Bio-One International) were treated with Cell-Tak solution (354240, Corning) to facilitate the cell adhesion. RPMs were seeded at 4 x 10^5^ cells per well in treated CELLview cell culture dishes. Seeded cells were incubated with or without 1 µg ml^−1^ of FITC-CTB (C1655, Sigma-Aldrich) for 12 h. Cells were then fixed with 4 % paraformaldehyde in PBS for 10 min, and blocked with 5 % normal goat serum (S-1000, Vector Laboratories Inc.). Dishes were stained with anti-IRE1α Ab (3294S, Cell Signaling Technology) or normal rabbit IgG (3900, Cell Signaling Technology) for Isotype control, and stained with Alexa546-Labelled Goat anti rabbit IgG (A11010, Invitrogen) and mounted with Vectashiled containing DAPI (H1200, Vector Laboratories Inc.). Stained dishes were analyzed with a confocal microscope, FV10i (Olympus).

## STATISTICAL ANALYSIS

Statistical significance was determined by an unpaired two-tailed Student’s *t*-test. *P* values are indicated by \**P* < 0.05, \*\**P* < 0.01, and \*\*\**P* < 0.001. *P* < 0.05 was considered statistically significant.

## Acknowledgements

We thank Ms. Aoi Matsumura-Tawaki, Ms. Akane Nishiwaki and Sumiko Okura for secretarial assistance, Ms. Ikuko Hattori and Chisako Mutsukawa for technical assistance. We acknowledge proofreading and editing by Benjamin Phillis at the Clinical Study Support Center, Wakayama Medical University.

## Author contributions

Conceptualization: I.S. and T.Kai. Methodology: S.M., H.H., T.T., K.H., S.F., and K.M. Investigation: I.S., Y.F.-O., C.N., N.W.-N., C.O., T.O., D.O., S.K., Y.F., A.Y., and T.Kat. Writing (original draft): I.S. and T.Kai. Writing (review and editing): I.S., S.M., H.H., T.T., K.H., and T.Kai. Resources: D.O., K.F., T.I., E.K. and K.J.I.

## Data and materials availability

RNAseq raw data have been deposited in the NCBI Gene Expression Omnibus database (GSE228915). All data needed to evaluate the conclusions in the paper are present or the Supplementary Materials.

## SUPPLEMENTARY MATERIALS

**Fig. S1.**
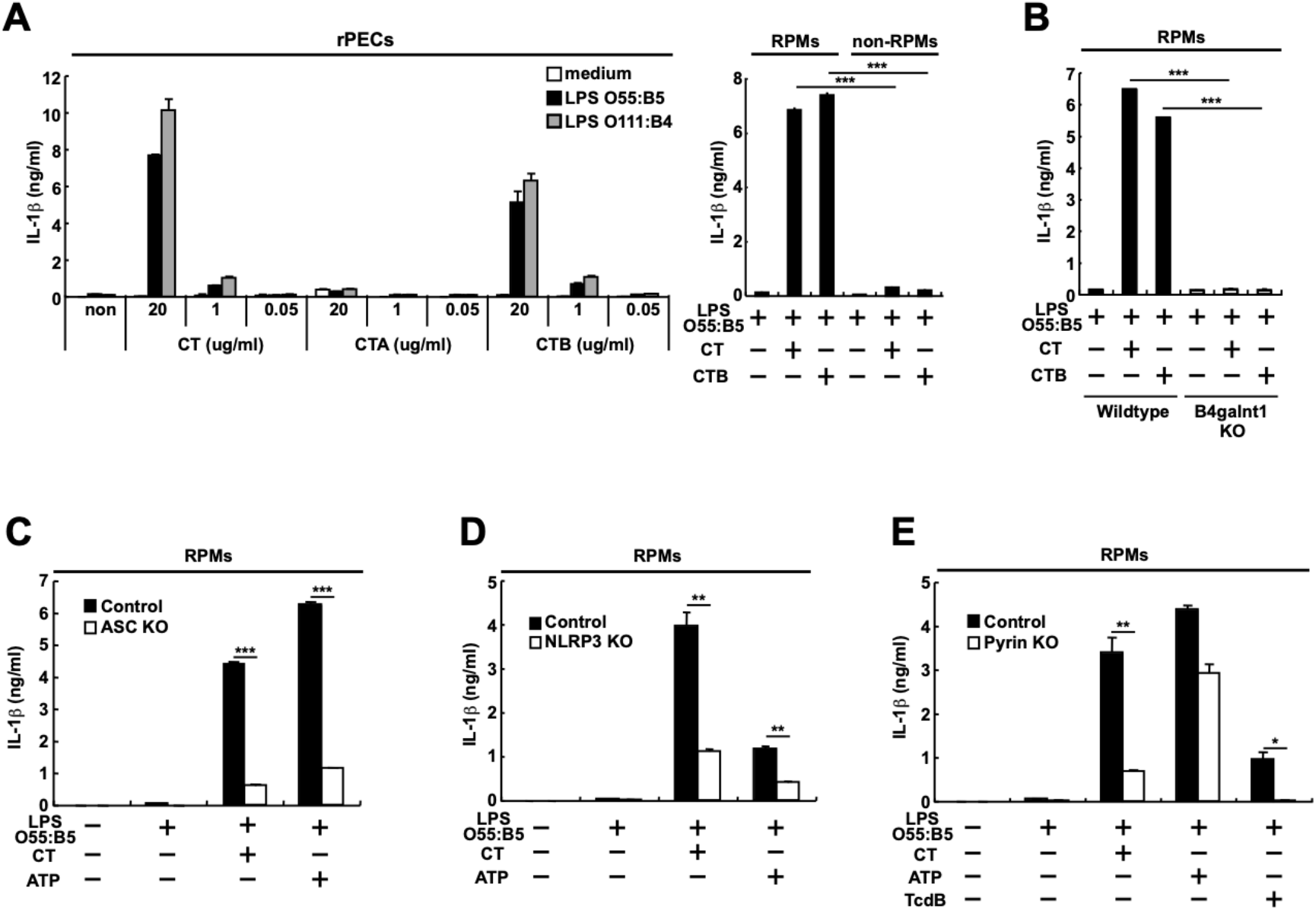
CT induces IL-1β production from LPS-primed RPMs through the pyrin inflammasome as well as the NLRP3 inflammasome. (A) On the left, rPECs from wild-type C57BL/6 mice were first cultured for 5 h in the absence (white bar) or presence of 500 ng ml^−1^ LPS O55:B5 (black bar) or LPS O111:B4 (gray bar). Then, 20 µg ml^−1^ CT, CTA or CTB were added and further cultured for 19 h. On the right, RPMs or non-RPMs from wild-type C57BL/6 mice were first cultured with 500 ng ml^−1^ LPS O55:B5 for 5 h. Then, 20 µg ml^−1^ CT or CTB were added and further cultured for 19 h. Culture supernatants were subjected to ELISA for IL-1β. The results are presented as means ± SD. \*\*\**P* < 0.001. Data are representative of two independent experiments. (B) RPMs from littermate B4galnt1 wild-type mice or B4galnt1-deficient mice were first cultured for 5 h in the presence of 500 ng ml^−1^ LPS O55:B5. Then 20 µg ml^−1^ CT or CTB were added and further cultured for 19 h. Culture supernatants were subjected to ELISA for IL-1β. Results are presented as means ± SD. \*\*\**P* < 0.001. Data are representative of four independent experiments. (C-E) RPMs from control or mutant mice lacking ASC (C), NLRP3 (D) or pyrin (E) were cultured for 5 h in the absence or presence of 500 ng ml^−1^ LPS O55:B5. Then, 20 µg ml^−1^ CT, 1mM ATP or 5 µg ml^−1^ TcdB (E) was added and further cultured for 19 h. Control mice were littermate ASC heterozygous mutant (C), littermate NLRP3 heterozygous mutant (D), or littermate pyrin wild-type mice (E). IL-1β production was measured by ELISA. Data are representative of three independent experiments. The results are presented as means ± SD. \**P* < 0.05, \*\**P* < 0.01, \*\*\**P* < 0.001.

**Fig. S2.**
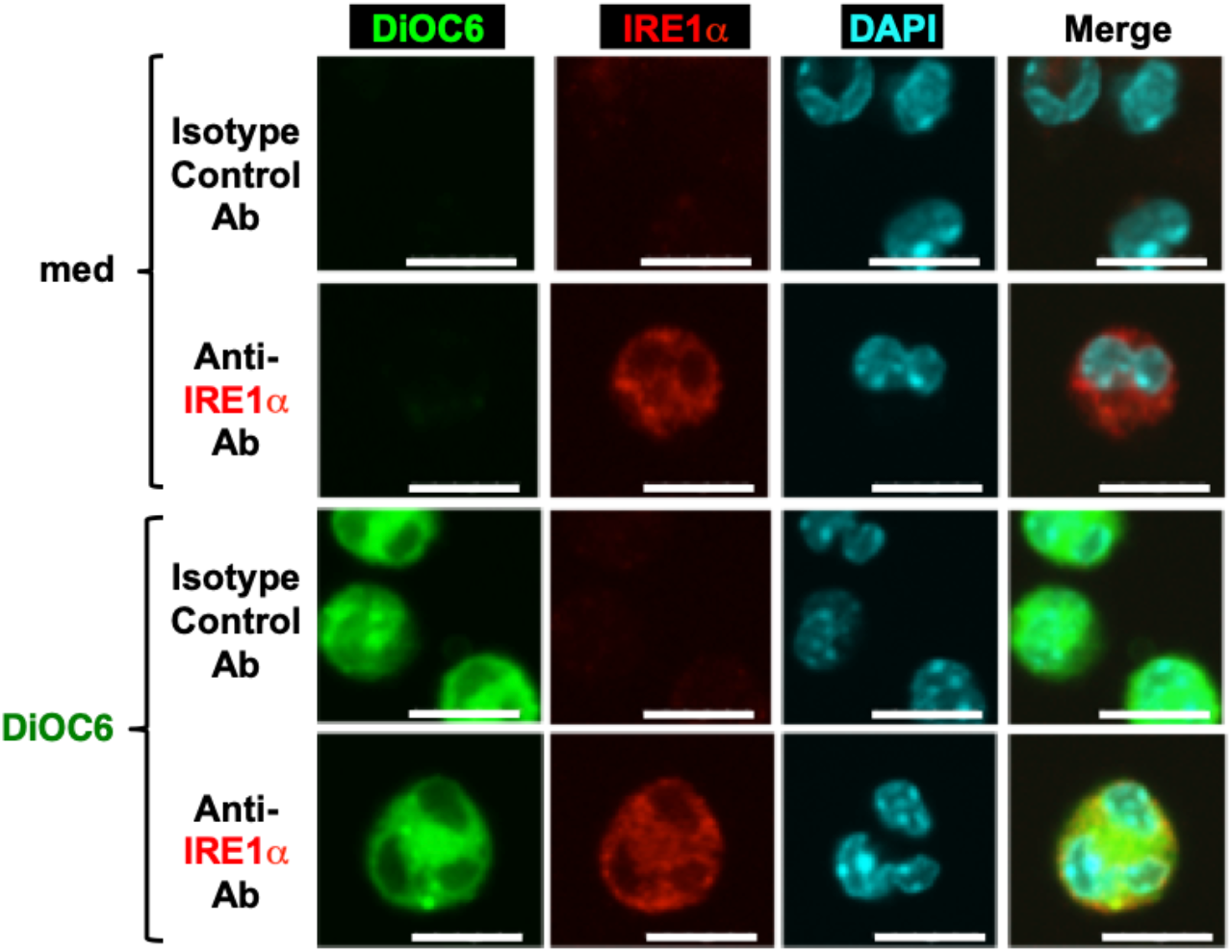
DiOC_6_ is co-localized with IRE1α in RPMs. RPMs from wild-type C57BL/6 mice were treated with or without 1µM DiOC_6_ (D-273, life technologies) for 5 min. Cells were then subjected to confocal microscopic analysis. Green, Red and Blue coloring represent DiOC_6_, IRE1α and DAPI staining, respectively. Rabbit IgG was used as Isotype control Ab. Scale bars represent 10 µm.

**Fig. S3.**
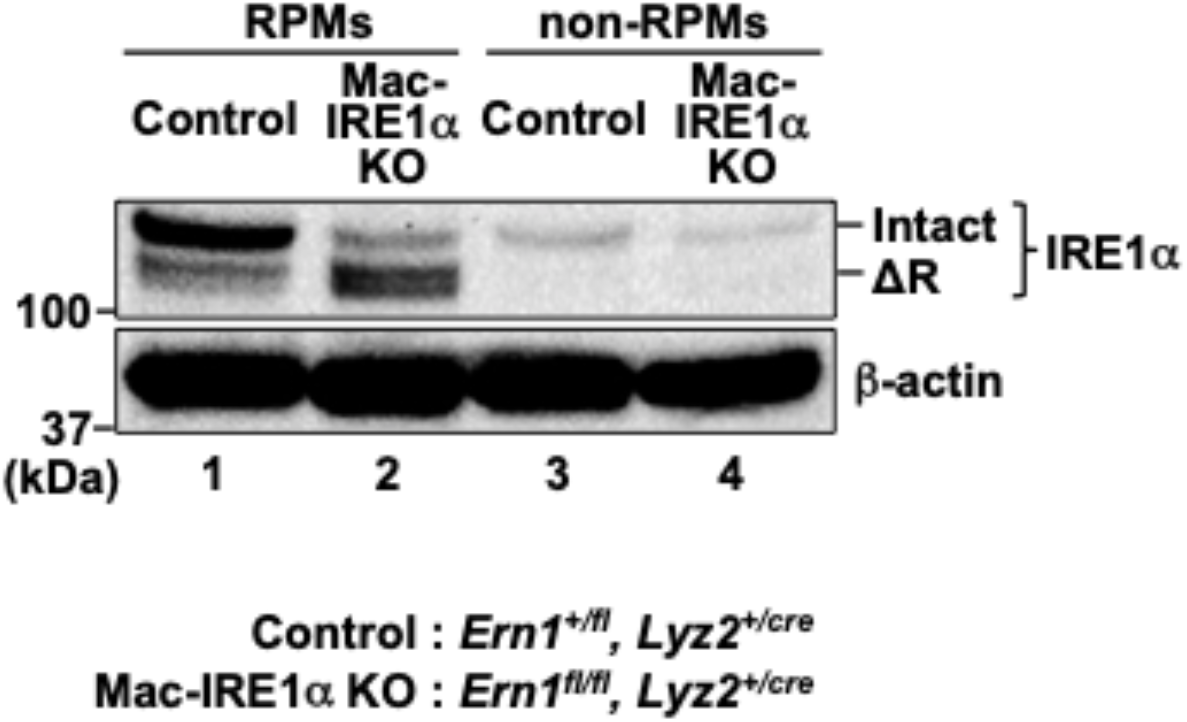
IRE1α-RNase domain is deleted in RPMs from Mac-IRE1α KO mice. Five hundred thousand cells of RPMs or non-RPMs cells from control or Mac-IRE1α KO mice were lysed in RIPA buffer with protease inhibitor cocktail. Lysate samples were then subjected to western blot for IRE1α. RNase domain-deleted IRE1α showed as ΔR. β-actin was used as a loading control.

**Fig. S4.**
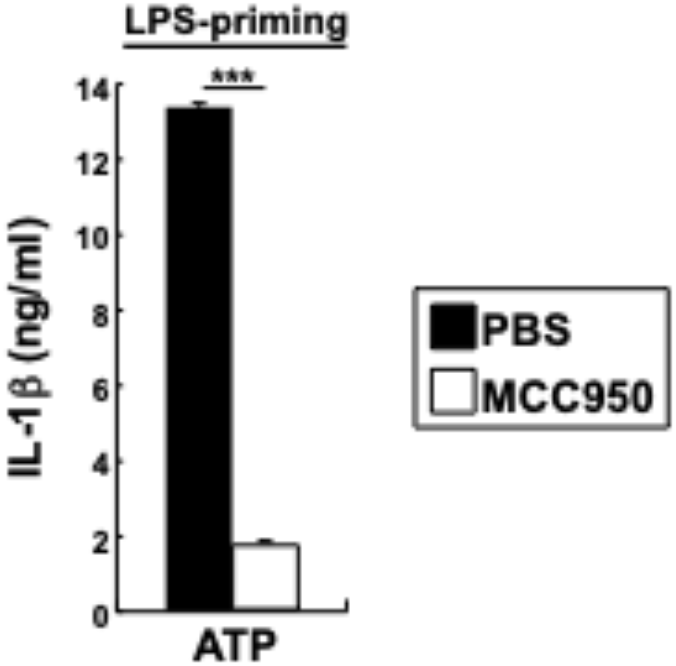
MCC950 inhibits ATP-induced IL-1β production from LPS-primed RPMs. RPMs from wild-type C57BL/6 mice were first cultured for 5 h in the presence of 500 ng ml^−1^ LPS O55:B5. Then 1 mM ATP was added and further cultured for 19 h. NLRP3 inhibitor, MCC950 (1 µM) or PBS as vehicle control were added 3 h before addition of ATP, TcdB. IL-1βproduction was measured by ELISA. The bars represent means ± SD. \*\*\**P* < 0.001.

